# Macromolecular toolbox to elucidate CLE-RLK binding, signaling and downstream effects

**DOI:** 10.1101/2024.01.11.570615

**Authors:** Madhumitha Narasimhan, Nina Jahnke, Felix Kallert, Elmehdi Bahafid, Franziska Böhmer, Laura Hartmann, Rüdiger Simon

## Abstract

Plant peptides communicate by binding to a large family of receptor-like kinases (RLKs) and they share a conserved binding mechanism, which may account for their promiscuous interaction with several RLKs. In order to understand the in vivo binding specificity of CLE peptide family, we have developed a novel set of CLAVATA 3 (CLV3) based peptide tools. After carefully evaluating the CLE peptide binding characteristics, using solid phase synthesis process, we have modified the CLV3 peptide and attached a fluorophore and a photoactivable side group. We observed that the labeled CLV3 shows binding specificity within CLAVATA1 clade of RLKs while avoiding the distantly-related PEP RECEPTOR clade, thus resolving the contradictory results obtained previously by many in vitro methods. Furthermore, we observed that the RLK-bound CLV3 undergoes clathrin-mediated endocytosis and gets trafficked to vacuole via ARA7-labeled endosomes. Additionally, modifying CLV3 for light-controlled activation enabled spatial and temporal control over CLE signalling. Hence our CLV3 macromolecular toolbox can be used to study rapid cell specific down-stream effects. Given the conserved binding properties, in the future our toolbox can also be used as a template to modify other CLE peptides.

**Highlight:** A macromolecular tool box consisting of modified CLE peptide with fluorescent molecule and photoactivable group offers reliable insights into its in vivo binding characteristics, localization and signaling.

## 1. Introduction

Small secreted peptides mediate cell-to-cell communication in multicellular organisms in response to pathogens, biotic and abiotic stimuli and to regulate developmental processes. The CLAVATA3 (CLE)/ EMBRYO SURROUNDING REGION (ESR)-RELATED (CLE) family of peptides is evolutionarily conserved among land plants. (Furumizu *et al*., 2021; Hirakawa, 2022; Whitewoods, 2021) CLE peptides act over short ranges to control stem cell division and differentiation in meristems, but also over long-distances between shoot and root tissues. The *Arabidopsis thaliana* genome comprises 32 *CLE* genes that encode 26 different CLE-peptides.(Hirakawa *et al*., 2011; Ito *et al*., 2006) CLE genes encode pre-pro-proteins containing an N-terminus signal peptide, a variable central domain which might be involved in the secretion process and a highly conserved CLE domain close to the C-terminus. After cleavage of signal peptide and proteolytic processing, the functional CLE-domain is composed of 12 or 13 amino acids that are secreted via ER and Golgi to the apoplast. The mature CLE-peptides often contain hydoxyproxiline residues that maybe essential for binding and activity, which can be further modified by arabinosylation.(Ito *et al*., 2006; Kim *et al*., 2017; Kondo *et al*., 2006; Rojo *et al*., 2002; Roman *et al*., 2022; Shinohara and Matsubayashi, 2013; Strabala *et al*., 2014; Tamaki *et al*., 2013; Xu *et al*., 2015)

The secreted peptide binds to the extracellular leucine-rich-repeat (LRR) domains of plasmamembrane (PM) localized receptor-like kinases (RLK), which interact with diverse co-receptors. Besides the LRR-ligand binding domain, the RLKs and their co-receptors carry a transmembrane domain and a cytosolic kinase domain that transmit the intracellular signal.(Shiu and Bleecker, 2001; Xi *et al*., 2019) The first identified CLE peptide is CLV3, which is expressed in the stem cell domain of the shoot apical meristem (SAM) of Arabidopsis. CLV3 signals via the LRR-RLK CLAVATA (CLV1), which belongs to the sub-family XI of LRR-RLKs (hereafter referred to as RLKs) that further includes BARELY ANY MERISTEM1 (BAM1), −2 and −3.(DeYoung *et al*., 2006; Furumizu *et al*., 2021) The CLV3 related peptide CLE40 is expressed in non-stem cells of root and shoot apical meristems (RAM and SAM, respectively) and signals via BAM1. During SAM development, the CLV3-CLV1 and the CLE40-BAM1 modules control expression of the transcription factor WUSCHEL, which is essential for meristem maintenance.(Schlegel *et al*., 2021; Schoof *et al*., 2000; Yadav *et al*., 2011) Overexpression or exogenous application of several different CLE peptides triggers premature differentiation of the SAM and root meristem. For example, both CLV3 and CLE40 peptides can induce arrest of root meristem growth and differentiation of root stem cells. This indicates promiscuity in the interaction between CLE peptides and RLKs.(Crook *et al*., 2020; DeYoung *et al*., 2006; Fiers *et al*., 2005; Hazak *et al*., 2017; Ito *et al*., 2006; Kinoshita *et al*., 2007; Narasimhan and Simon, 2022) This has been corroborated by in vitro studies on isolated receptor domains in which dissociation constants between 0.6nM and 14µM (K_d_) were observed for CLE peptides and RLKs of subfamily XI. For example, CLE8-14 and −16 can bind BAM1 with very high affinities, but CLE9 can also bind CLV1.(Crook *et al*., 2020; Ogawa *et al*., 2008) Notably, CLV3 has been shown to bind to the RLKs CLV1, BAM1, −2 and −3, which belong to the CLV1 clade.(Crook *et al*., 2020; Furumizu *et al*., 2021; Guo *et al*., 2010; Hazak *et al*., 2017; Shinohara *et al*., 2012)

Interestingly, some CLEs were found to interact with RLKs of different clades. CLE9, in addition to BAM1, binds also to HAESA-LIKE1 (HSL1) with lower affinity. HSL1 is an RLK that predominantly interacts with the peptide INFLORESCENCE-DEFICIENT IN ABSCISSION (IDA) and other members of the IDA-LIKE family of ligands(Furumizu *et al*., 2021; Qian *et al*., 2018; Roman *et al*., 2022). CLE14, in addition to BAM1, was also reported to interact with PEP RECEPTOR 2 (PEPR2), which is a receptor for the AtPep family of peptides.(Gutierrez-Alanis *et al*., 2017) This broad interaction spectrum can be attributed to the conserved mode of binding, which is shared amongst CLEs and other peptide families. The C-terminus residues undergo close contacts with the RLK and their coreceptors.(Li *et al*., 2017; Zhang *et al*., 2016a; Zhang *et al*., 2016b) The N-terminal residues would then interact only with the RLK and provide specificity for the interaction. For example, changing the first three N-terminus residues of the CLE peptide TRACHEARY ELEMENT DIFFERENTIATION INHIBITORY FACTOR (TDIF or CLE41) or of CLE9 resulted in total loss of binding to their cognate RLKs. Furthermore, swapping one or more of the N-terminus residues between IDA and CLE9 resulted in swapped affinity to their cognate RLKs HSL1 and BAM1, respectively.(Li *et al*., 2017; Roman *et al*., 2022; Zhang *et al*., 2016a) Thus, N- and C-termini form two anchoring sites for the peptide on the RLKs, with the N-terminus contributing to the specificity of interactions. Furthermore, the peptides flg22, derived from the flagellin protein, and pep1 have been shown to bind FLAGELLIN-SENSITIVE2 (FLS2) and PEPR, respectively, in a fully extended conformation, which is in stark contrast to CLE41 forming a kink-like structure in the middle while binding the RLK TDR.(Zhang *et al*., 2016b) This indicates that peptides of different families have mechanisms that confer specificity to the RLKs of a certain clade, thus excluding unspecific RLK interactions. This brings the previously reported CLE14-PEPR2 interaction into question, particularly when PEPR clade is phylogenetically distant from the CLV1 and TDR clades, which act as CLE receptors. (Furumizu *et al*., 2021; Gutierrez-Alanis *et al*., 2017) Two important questions remain unanswered: Can CLE peptides interact with high affinity and induce signaling through the PEPR clade of receptors? Is there an in vivo binding affinity between CLEs and specific RLK clades, such as CLV1 and TDR sister clades, while eliminating cross-clade interaction with phylogenetically more distant RLKs?

Most of our understanding on peptide-receptor interaction comes from in silico models, in vitro studies with purified RLKs in stable conditions, or in heterologous system.(Gutierrez-Alanis *et al*., 2017; Ito *et al*., 2006; Li *et al*., 2017; Ogawa *et al*., 2008; Roman *et al*., 2022; Shinohara *et al*., 2012; Zhang *et al*., 2016b) However, such studies do not accurately reflect the in vivo conditions, such as receptor confirmation changes within multi-protein complexes, the chemical and physical environment, protein modifications, number of receptor molecules at the PM, ligand diffusion kinetics etc.(Bhattacharya *et al*., 2013; Chang, 2022; Kastritis and Bonvin, 2013).

Post CLE peptide interaction, the fates of the peptide and the RLK have not been extensively studied. In plants, several receptors have been shown to be internalized via clathrin-mediated endocytosis (CME).(Paez Valencia *et al*., 2016) Different RLKs and ligands have been shown to undergo CME with varying rates and subsequent endosomal trafficking. For example, pep1-PEPR complexes undergo a slower rate of CME compared to BRASSINOSTEROID-INSENSITIVE1 (BRI1). This could be attributed to the pre-formed BRI1-co-receptors complexes at the PM to receive brassinosteroid.(Bucherl *et al*., 2013; Ortiz-Morea *et al*., 2016) Similarly, CLV3 peptide binding can induce rapid formation of larger multimeric aggregates of CLV1 with co-receptors at the PM. In the SAM, CLV1 is endocytosed and trafficked towards the vacuole for degradation.(Nimchuk *et al*., 2011; Somssich *et al*., 2015; Wang *et al*., 2023) However, the sub-cellular trafficking dynamics of CLE peptides and their fate within the cell after RLK binding are unknown.

To enable in vivo studies of CLE-receptor interactions, we developed two new tools: (i) we synthesized a functional, fluorescently labeled CLV3 peptide in order to track the spatial and temporal distribution of CLE peptides and their cognate RLKs in vivo, and (ii) generated a photocaged CLV3 peptide that can be activated by light with high spatial and temporal control. Fluorophore labeled pep1, flg22 and brassinosteroid hormone have been used previously to understand their cognate receptor-mediated signaling and subsequent sub-cellular dynamics.(!!! INVALID CITATION !!! 37,45,46) Based on the existing knowledge on CLE-RLK recognition mechanism and key interacting residues(Yamaguchi *et al*., 2016), we modified the CLE motif of the CLV3 peptide to allow linkage of a fluorescent dye, while retaining specific binding activity to the RLKs. This was achieved by conjugation of a fluorophore to the side chain. To accomplish this, we substitute the second amino acid threonine, previously identified as non-essential for binding in depletion assays(Fiers *et al*., 2006), to lysine and linked a suitable fluorophore to this position. A quantum-dot based probe for CLV3 was previously reported, but it could not be used in planta due to toxic effects.(Yu *et al*., 2014) Our functional CLE peptide probe allows to study in vivo binding specificity and subsequent sub-cellular trafficking.

We then used photo-caging to obtain a peptide probe that can be activated locally and rapidly using light (Mangubat-Medina and Ball, 2021). A photocage is a photolabile protecting group that can be used to block the functional groups of the peptide required for receptor binding and biological activity. The caged, non-binding peptide can then be transformed into its active form by releasing the cage group through sample illumination with a specific wavelength of light (Figure 1). So far, three molecular plant hormones - auxin, abscisic acid, and gibellerin - have been studied with the aid of photocaging techniques.**(Kusaka *et al*., 2009)** However, no photoactivatable and fluorescently labeled plant peptide has been synthesized and applied in planta so far.

**Figure 1.**
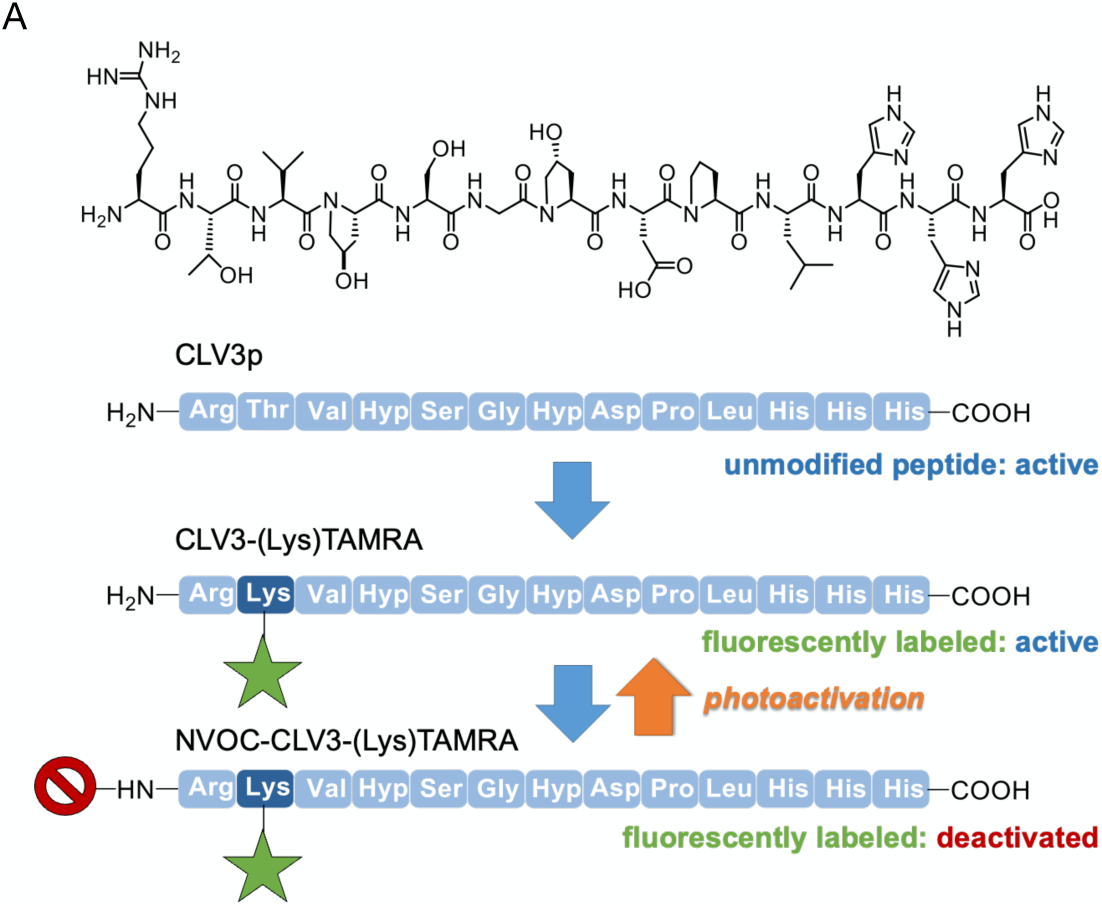
CLV3 sequence and schematic presentation of the modifications. A) Introduction of a fluorophore (green) to CLV3to derive an active CLV3 probe followed by introduction of a N-terminal photocage (red) to allow for light activated binding to CLV1 receptor in planta.

In this paper, we have presented the development of a macromolecular tool box of CLE peptides including fluorescently-labeled and photoactivatable peptides. Performing a series of in vivo tests, we have demonstrated their bioactivity, specificity and ability to be locally activated. Using these tools, we have further demonstrated the binding specificities and subcellular dynamics in a spatially and temporally controlled manner. Overall, we have provided a blue print of how to derive a truly functional and controllable CLE peptide to understand the recognition and binding capacity to its cognate RLKs.

## 2. Results

### 2.1 Synthesis of a functional, fluorescently labeled CLV3 probe (CLV3-TAMRA)

Arabidopsis CLE domains share several conserved residues (Figure 2A) indicating that they interact with RLKs through a conserved mechanism.(Zhang *et al*., 2016a; Zhang *et al*., 2016b) We chose CLV3, one of the best understood CLEs, as a test system.

**Figure 2.**
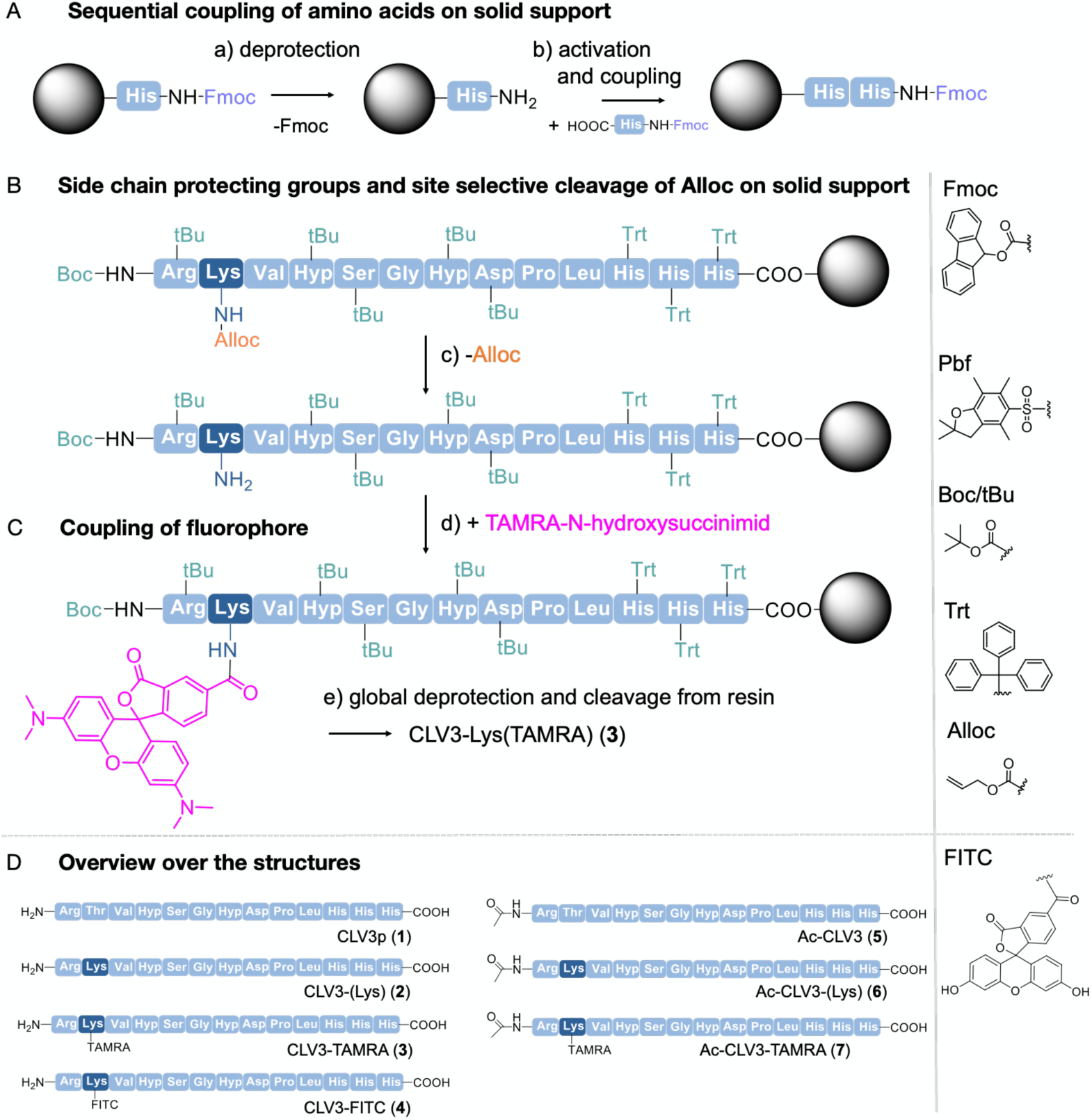
Solid phase peptide synthesis of CLV3 using preloaded Fmoc-His(Trt)-Resin and Fmoc-protecting group (PG) standard protocol. A) Deprotection of N-Terminal Fmoc-PG (a) using 25 Vol% piperidin in DMF for 5-, 15- and 20min. Coupling of Fmoc-protection amino acid (AA) (b) using 5 eq. AA, 5eq. PyBOP and 10 eq. DIPEA in DMF for 1h. Repeating both steps a)/b) to build up sequence. B) Selective alloc deprotection (c) of lysin under reductive conditions with tetrakis(triphenylphosphine) palladium (0) catalyst and 10 eq. 1,3 dimethylbarbituric acid in DCM two times for 45 min. C) Conjugation of TAMRA-N-hydroxysuccinimid (d) at free Lysine by adding 10 eq. DIPEA in DMF for 16h. Cleavage of the final structure (e) with 70 Vol% TFA and 30 Vol% TIPS. D) Overview of different structures. See Supplemental File 1 for further details on the synthesis and analytical data of different peptide probes.

Synthesis of all modified peptides was performed on an automated peptide synthesizer using well-established solid phase peptide synthesis employing fluorenylmethyloxycarbonyl (Fmoc) protected amino acids.(Wellings and Atherton, 1997) In short, the terminal carboxy group of the amino acid is activated *in situ* to form an active ester that allows for coupling to a resin-bound amine group e.g., the N-terminus of the previous amino acid, at room temperature and with high yields. Upon coupling of the amino acid, the Fmoc group is selectively cleaved by piperidin, releasing the N-terminus now available for coupling of the next amino acid. After successful build-up of the desired amino acid sequence through such iterative coupling, the peptide is cleaved from the resin, typically under acidic conditions, including the cleavage of potential side chain protecting groups. Synthetic protocols for the unmodified CLV3 peptide were developed by choosing a suitable side chain protecting group strategy (Figure 2A). (2,2,4,6,7-pentamethyldihydrobenzofuran-5-sulfonyl(Pbf) for arginine, tert-butyloxycarbonyl (tBu) for hydroxyproline, serine and aspartic acid and trityl (trt) for histidine). Upon final cleavage of the peptide from the resin using acidic conditions, the fully deprotected CLV3 peptide (CLV3p) (**1**) was isolated in high purity (Data S1, Data S2 and Data S3). Based on this protocol, a series of modified and fluorescently labeled CLV3 peptides was synthesized (Figure 2D). One of the challenges in modifying small peptide probes with fluorescent labels is the decrease or even loss of their biological activity e.g., due to a change in polarity, capping of essential amino acid residues or sterical shielding of peptide sites that are required for interaction with the receptor.(Boaro *et al*., 2020) Previous studies showed that the C-terminal residues interact with the receptor-coreceptor interface in a conserved manner, and that the N-terminal residue, either R or H, undergoes specific interactions with the receptor. The first and third residues are vital for recognizing and anchoring the CLE peptide to the RLK. The second residue is less conserved and can be modified without strongly compromising peptide activity.(Li *et al*., 2017; Song *et al*., 2012; Zhang *et al*., 2016a) Therefore, the threonine at position 2 in CLV3p was replaced by lysine, giving CLV3-(Lys) (**2**). The amino side chain of the lysine amino acid allows for the introduction of another orthogonal protecting group, allyloxycarbonyl (Alloc), which can be cleaved under reductive conditions on a solid support (Figure 2B).(Wojcik *et al*., 2012) Therefore, the N-terminal amino acid arginine was changed from Fmoc to a Boc protecting group, as during reductive deprotection of the lysine side chain partial deprotection of the N-terminal Fmoc protecting groups was observed. Selective release of Alloc allowed for quantitative and site-selective introduction of a fluorophore, here either TAMRA or fluorescein (FITC), giving the two fluorescently labeled CLV3 probes CLV3-TAMRA (**3**) and CLV3-FITC (**4**) (Figure 2C,D). TAMRA and fluorescein were chosen as they are both commonly used in fluorescence microscopy, but differ in their excitation and emission spectra (λex,max = 550nm and λem,max = 580nm for TAMRA and λex,max = 500nm and λem,max = 520nm for FITC).(Tung, 2004)

In order to assess the binding characteristics of CLV3-TAMRA as a peptide probe, we synthesized negative controls, a series of CLV3 peptides that are acetylated at the N-terminus (Figure 2D). Instead of the Boc-protected arginine, its Fmoc-protected variant was used. Fmoc was cleaved off under basic conditions while every other protection group was stable under these conditions and capped with acetic anhydride to receive the acetylated N-Terminus.

### 2.2 Testing the bioactivity of CLV3-TAMRA

Bioactivity of the modified peptides was tested using a root length assay. Exogenous application of the synthesized native CLV3 peptide (CLV3p) elicits a premature differentiation of the root meristem, leading to a short root phenotype, at CLV3p concentrations of 10nM or higher.(Blümke *et al*., 2021; Hazak *et al*., 2017) We compared root lengths of seedlings grown on medium with different concentrations of CLV3p or the modified peptides. Seedlings showed a strong reduction root length in response to CLV3p at 100nM; CLV3-TAMRA was ineffective at 100nM, but triggered root length reduction at 1uM concentration. Neither Ac-CLV3-TAMRA, free TAMRA fluorophore or CLV3-FITC elicited any response at 1uM (Figure 3, S1). Signalling of CLV3p during root development depends on the CLV2/CRN-receptor heteromer.(Fiers *et al*., 2005; Miwa *et al*., 2008) Arabidopsis seedlings mutant for CRN (*crn*) did not respond to 1uM a concentration of the tested peptides, indicating that the CLV3-TAMRA peptide acts through the canonical CLV3 signaling pathway, although with a lower efficacy than the unmodified CLV3p (Figure 3, S1A). Lack of CLV3-FITC biological activity could be due to structural differences between the fluorophores: TAMRA carries tertiary amines in the xanthene core and is unable to form hydrogen bonds, while FITC contains hydroxyl groups that could interact with the peptide backbone, potentially altering its conformation and preventing binding to the receptor protein. In summary, modifying the second amino acid residue of the CLE domain to attach the fluorophore is an effective strategy in creating a bioactive CLE peptide probe. TAMRA, as a fluorescing agent for CLE peptide probes, can retain their bioactivity, although with a slightly diminished efficiency.

**Figure 3:**
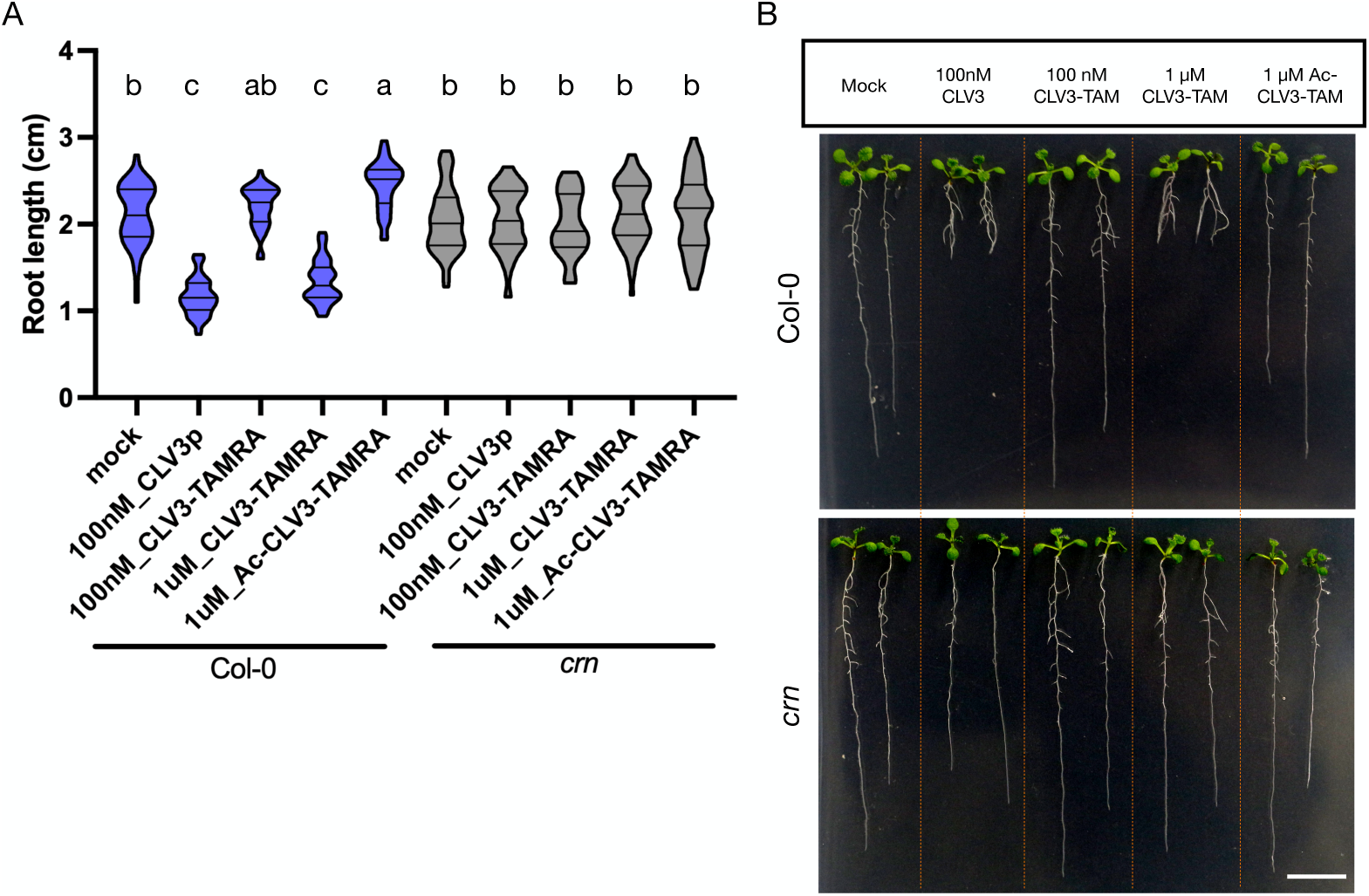
Bioactivity of CLV3-TAMRA. A) Violin plot representing root length analyses of Col-0 and *crn* after 7-day treatment with mock, CLV3p, CLV3-TAMRA and Ac-CLV3-TAMRA. n≥ 43 roots each condition; N=3. The lines represent the median and the quartiles. Two-way ANOVA with interaction, F test p<2e-16 ***. Statistical grouping was calculated by Tukey’s HSD test (**α**= 0.001). Groups sharing the same letter are not significantly different. B) Images of seedlings of Col-0 and *crn* after 10-day treatment. Scale bar: B – 1 cm.

### 2.3 Sub-cellular localization and trafficking of CLV3-TAMRA

We tested if CLV3-TAMRA is recognized by the RLKs at the PM, and how it interacts with the endocytic sub-cellular trafficking machinery. When added to Arabidopsis root or shoot meristems, CLV3-TAMRA localized to the PM in all layers of the meristems within 3min after addition (Figure 4A, B, S2A). The negative control peptide Ac-CLV3-TAMRA showed a very weak signal at the PM, possibly due to unspecific binding to other PM proteins (Figure 4A, B).

**Figure 4:**
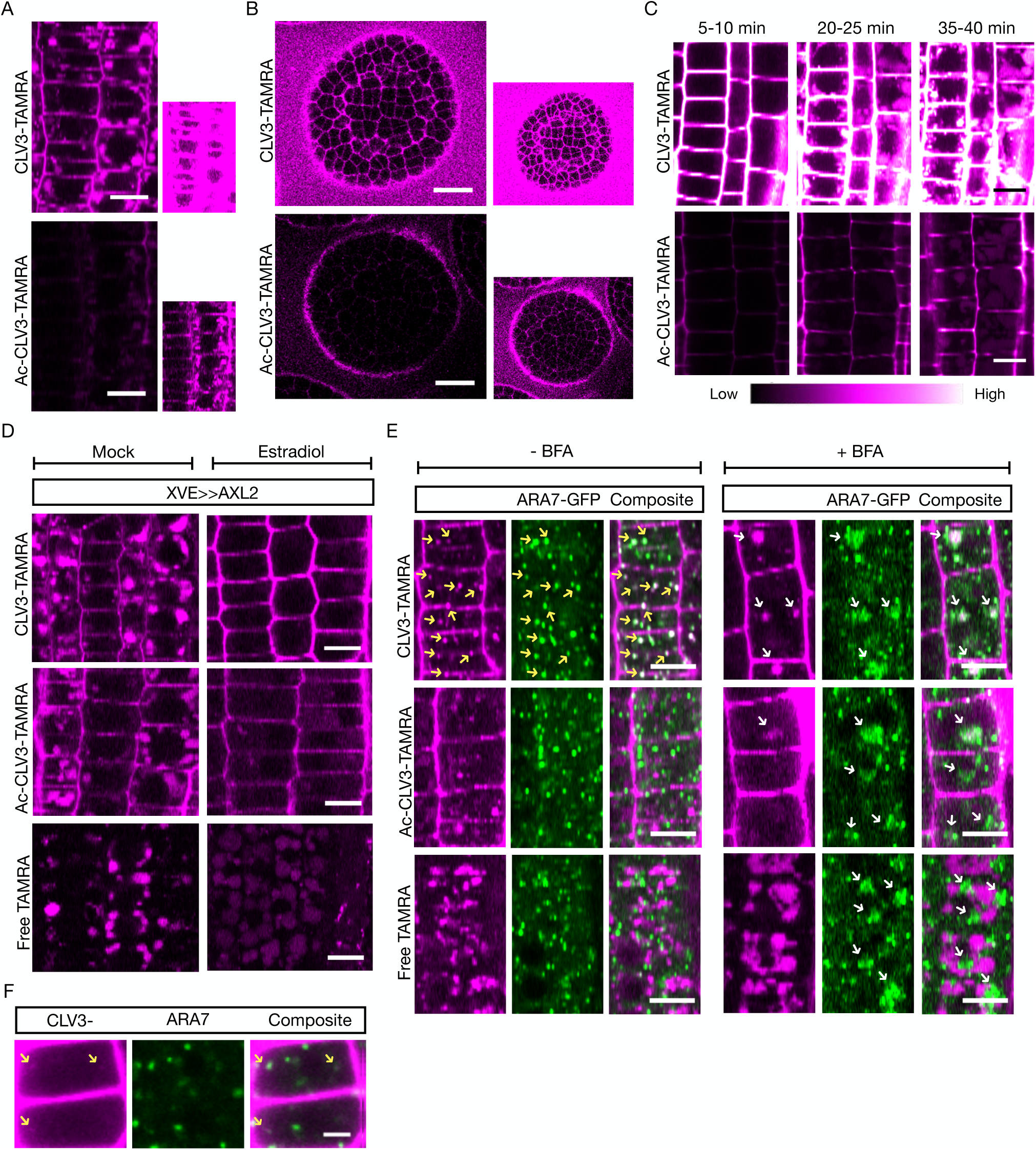
Sub-cellular localization and trafficking of CLV3-TAMRA. A-F) Representative confocal images of root meristem epidermal cells (A, C, D-F) or SAM cells (B) after 1 µM treatment of the peptide CLV3-TAMRA, or the controls Ac-CLV3-TAMRA or free TAMRA fluorophore. Lines used: A and B – Col-0; C – *pBAM1::BAM1-GFP; bam1-3*; D – *XVE>> AUXILIN-LIKE2*; E and F – *p35s::ARA7-GFP.* A) PM localization after 15 min treatment. n=6; N=2. The inset on the right shows the same images with increased brightness. B) PM localization in SAM cells after 1 min treatment and washing. N≥5. The inset on the right shows the same images with increased brightness. C) Vacuolar accumulation after continuous treatment overtime. n≥6; N=2. D) Clathrin-mediated endocytosis and vacuolar trafficking after 30 min treatment. The seedlings either had induction of AUXILIN-LIKE 2 overexpression under estradiol treatment or no induction under mock treatment. n≥6; N=2. E) Localization of peptide and controls in LE 10 to 15 min post 5 min treatment. n≥5; N=2 (left); in BFA bodies after 50 min treatment. n≥5; N=3 (right). F) Localization of the peptide in LE after 4 min incubation. n=4; N=2. Yellow arrows indicate the CLV3-TAMRA co-localised with LEs marked by ARA7-GFP. White arrows indicate the BFA bodies containing LEs, and the peptide or the controls. Note: In D-F, Ac-CLV3-TAMRA and Free TAMRA were imaged at higher laser intensity and gain than CLV3-TAMRA to observe their sub-cellular localization. Scale bar: A, C-E – 10 µm; B – 15 µm; F – 3 µm.

During the course of 40min treatment, CLV3-TAMRA accumulated increasingly in the vacuole, which is surrounded by the tonoplast (Figure 4C, S2B). We first tested if the accumulating vacuolar signal is due to endocytosed CLV3-TAMRA peptide, or free TAMRA fluorophore that resulted from CLV3-TAMRA instability or degradation. We analyzed CLV3-TAMRA from our stock solution via RP-HPLC after 10 months storage, and saw no degradation of the fluorophore, which indicates that the TAMRA fluorophore remains stably bound to the CLV3-peptide (Data S4). We concluded that CLV3-TAMRA is functional and reaches the vacuole through the endocytic sub-cellular trafficking pathway.

We then examined the CLE peptide sub-cellular trafficking process. In animal cells, after endocytosis, ligand-bound receptors enter the early endosome for sorting. The membrane bound receptors are primarily recycled and return to the PM, but ligands are mostly delivered to the late endosome (LE) and then degraded.(French and Lauffenburger, 1997; Lakadamyali *et al*., 2006; Solinger and Spang, 2022) We first examined if CLV3-TAMRA undergoes CME from the PM using AUXILIN-LIKE2 over-expression, which strongly hinders CME.(Adamowski *et al*., 2018) After 60min of CLV3-TAMRA treatment, CLV3-TAMRA accumulated in the vacuole. However, estradiol inducible over-expression of AUXILIN-LIKE2 strongly diminished the vacuolar signal compared to mock treatment, showing that CLV3-TAMRA is subject to CME (Figure 4D). The inactive Ac-CLV3-TAMRA and free TAMRA fluorophore also reached the vacuole (Figure S2B), although to a lesser extent. CME inhibition interfered with uptake and vacuolar localization of Ac-CLV3-TAMRA (Figure 4D), indicating that it might still bind PM localized RLKs with reduced affinity, possibly via the conserved C-terminal interaction. Free TAMRA fluorophore was neither bound at the PM nor trafficked via endosomes; therefore, it likely entered cells via diffusion (Figure S2C).

We then investigated the endocytic route to the vacuole. We found that CLV3-TAMRA co-localized with a few ARA7-labeled LEs(Lee *et al*., 2004) rapidly within 5 min (Figure 4E - left, 3F). After 15 minutes, CLV3-TAMRA reached almost all the ARA7-labeled LE compartments and aggregated into BFA-bodies after Brefeldin A (BFA) treatment (Geldner *et al*., 2009; Narasimhan *et al*., 2021) (Figure 4E - right). However, we only occasionally observed localization of CLV3-TAMRA in early endosomes (EEs) (Figure S2D). This is in concordance with flg22 and pep1 ligands that are also predominantly localized in LEs en route to vacuole,(Jelenska *et al*., 2017; Ortiz-Morea *et al*., 2016) and in contrast to animal cells where the ligand-receptor sorting occurs in EEs. However, neither free TAMRA nor Ac-CLV3-TAMRA co-localized with LEs (Figure 4E - left), although a weak Ac-CLV3-TAMRA signal was observed in BFA bodies (Figure 4E - right). We conclude that CLV3-Tamra is endocytosed via CME and effectively trafficked along the endosomal pathway. When and where the separation of endocytosed peptide ligands from the RLKs occur in plant cells remains to be solved. Overall, we found that CLV3-TAMRA endocytosis and sub-cellular trafficking dynamics is very rapid, and that the trafficking route is concordant with other plant signaling peptides.

### 2.4 Binding specificity of CLV3-TAMRA to RLKs

Studies in heterologous system reported that CLV3 binds to the RLKs CLV1 and BAM1, although reports on binding affinities vary significantly between studies.(Crook *et al*., 2020; Guo *et al*., 2010; Hazak *et al*., 2017; Ogawa *et al*., 2008) With our probes in hand, we here tested the interaction of CLV3-TAMRA to CLV1 and BAM1 in vivo.

After CLV3 binding, CLV1 in the SAM undergoes endocytosis and is targeted to the vacuole. The CLV1 population at the PM is strongly reduced over time.(Nimchuk *et al*., 2011; Wang *et al*., 2023) We observed that CLV3-TAMRA, but not Ac-CLV3-TAMRA, induced CLV1 endocytosis and vacuolar targeting (Figure 5A). However, CLV3 binding does not trigger endocytic loss of the BAM1 PM pool, or increased vacuolar trafficking of BAM1 (Figure S3A). This difference could be due to a high rate of recycling of BAM1, or alternatively due to a smaller fraction of BAM1 being involved in active signaling that further undergo degradation.(Bucherl *et al*., 2013; Sorkin and von Zastrow, 2009) Therefore, to test for interaction with BAM1, we used a transgenic Arabidopsis line that overexpressed BAM1-GFP in the root meristem (Figure S3B). Compared to wild-type root meristems, significantly higher amount of CLV3-TAMRA bound to the PM of BAM1-GFP (OX) root meristems than to wildtype (Figure 5B, S3C). PM binding of Ac-CLV3-TAMRA was generally very low, but slightly increased in the BAM1-GFP (OX) roots (Figure S3C). On the other hand, we observed a decrease in PM bound CLV3-TAMRA in the stele layers of *bam1* and *bam1;bam2* mutants compared to WT, although there was no obvious difference in the epidermal layer. It is to be noted that this decrease could be attributed to an increased endocytic rate in the stele of the mutants as indicated by strong cytosolic signal (Figure S3D). We then incubated root meristems of BAM1-GFP (OX) line with CLV3p and CLV3-TAMRA to test if both compete for the same binding sites on the PM (Figure 5C). With increasing concentration of CLV3p in the medium, the amount of CLV3-TAMRA fluorescence on the PM decreased, showing that CLV3p can compete with CLV3-TAMRA binding to the same site. This demonstrates that CLV3-TAMRA has strong affinity to both CLV1 and BAM1 in vivo, in contrast to results obtained from in vitro studies.

**Figure 5:**
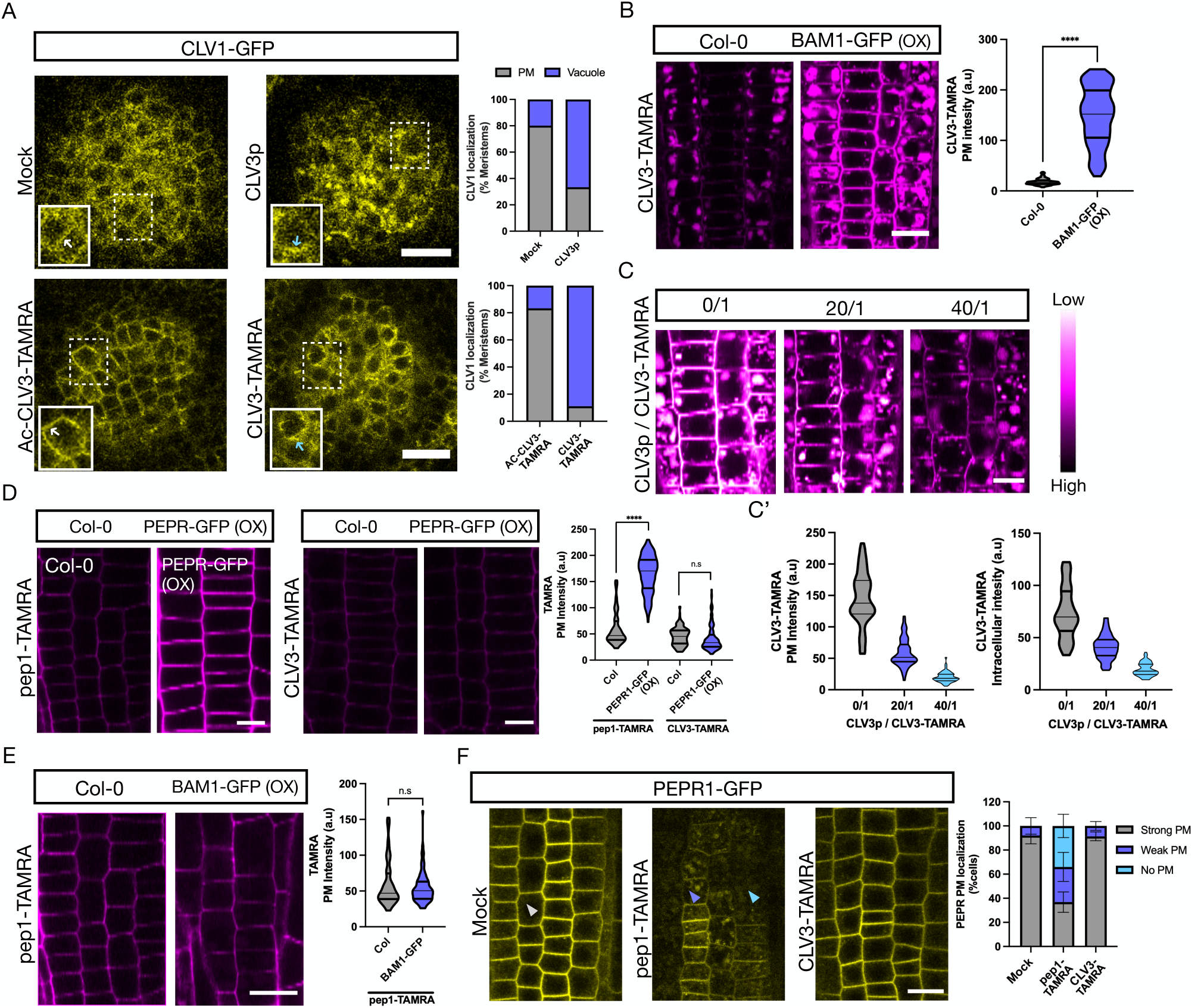
Binding specificity of CLV3-TAMRA to RLKs. A-F) Representative confocal images of SAM (A) or root meristem epidermal cells (B-E) after respective treatments. Lines used: A – *pCLV1::CLV1-2X-GFP*; *clv1-11;* B and E – Col-0; *pBAM1::BAM1-GFP*;*bam1-3;* C – *pBAM1::BAM1-GFP*; *bam1-3;* D – Col-0; *pPEPR1::PEPR1-GFP*; *pepr1; pepr2*; F – *pPEPR1::PEPR1-GFP*; *pepr1; pepr2*. A) CLV1 localization after 30 min of treatment with either mock or 100 nM CLV3 (top). N≥5; 1 µM of either Ac-CLV3-TAMRA or CLV3-TAMRA (bottom). N≥9. The insets showcase the PM (white arrow) or vacuolar localized CLV1 (blue arrow) in a cell. Stacked bar graph represents the relative number of SAMs with PM and vacuolar localized CLV1. B) CLV3-TAMRA PM signal after 1 µM, 30 min treatment in Col-0 or BAM1-GFP over-expression line. n≥6; N=3. The violin plot represents CLV3-TAMRA mean fluorescence intensity of the PM from 13-39 cells/root. Two-sided t test. For Col-0 > BAM1-GFP p<0.0001. C) CLV3-TAMRA PM signal after 30 min treatment with CLV3p/CLV3-TAMRA combination – (from left to right) 0 µM/1 µM, 20 µM/1 µM, 40 µM/1 µM. n≥7; N=2. C’) The violin plots represent the mean fluorescence intensity of the PM (top) or the intracellular region (bottom) from 5-17 cells/root. D) pep1-TAMRA and CLV3-TAMRA PM signal after 200nM, 1 min treatment in Col-0 and PEPR1-GFP over-expression line. n=6; N=1. The violon plot represents TAMRA mean fluorescence intensity of the PM from 14-31 cells/root. Two-sided t test. For pep1-TAM/PEPR-GFP > Col-0 p<0.0001; For CLV3-TAM/PEPR-GFP > Col-0 p=n.s. E) pep1-TAMRA PM signal after 200 nM, 1 min treatment in Col-0 or BAM1-GFP over-expression line. n≥6; N=1. The violon plot represents pep1-TAMRA mean fluorescence intensity of the PM from 10-25 cells/root. Two-sided t test. For BAM1-GFP > Col-0 p=n.s. F) PEPR1 PM localization after 30 s pulse with mock, 100 nM pep1-TAMRA or 1 µM CLV3-TAMRA. Grey, violet and blue arrow heads indicate an example cell with strong, weak and no PM PEPR. The stacked bar graph represents relative number of cells with strong, weak or no PEPR at the PM. n=5; N=2. At least 55 cells were analyzed/root. Error bars indicate mean ± SD. The lines in the violin plots indicate the median and the quartiles. Scale bar: A, B, E, F – 15 µm; C, D – 10 µm.

Because CLV3 has a strong affinity to CLV1 clade RLKs, we tested for its cross-clade interaction with PEPR1, an RLK of a phylogenetically distant clade.(Furumizu *et al*., 2021) pep1 and pep1-TAMRA bound and induced endocytosis of its PM localized cognate receptor PEPR1-GFP, causing a significant downregulation of the PM PEPR population through vacuolar targeting(Ortiz-Morea *et al*., 2016) (Figure 5D, F, S3E). However, CLV3-TAMRA did not exhibit an increased binding in a PEPR1 over-expression transgenic line (Figure 5D); moreover, neither CLV3p nor CLV3-TAMRA induced PEPR endocytosis from the PM (Figure 5F, S3E). In addition, pep1-TAMRA, did not show increased binding affinity to BAM1 (Figure 5E). We conclude from our in vivo studies with modified CLV3 peptides that interactions of peptides are limited to a specific RLK family, pep1 to the PEPR clade and CLV3 to the CLV1 clade, contradicting earlier reports of cross-clade promiscuity in CLE peptide interactions.(Furumizu *et al*., 2021; Gutierrez-Alanis *et al*., 2017; Qian *et al*., 2018)

### 2.5 Spatial and temporal control of CLE signaling

CLV3-TAMRA is a functional bioactive probe that, when externally applied, will rapidly reach all tissue layers and elicit a response. In order to achieve a controlled activity of the peptide in the targeted tissue or at a specific time point, we used nitroveratryloxycarbonyl (NVOC) as photo-cleavable protecting group, or photocage for the N-terminus of the CLV3 peptide with a maximum absorption for cleavage at 365nm.(Kneuttinger, 2022) Linking the NVOC group to the N-terminus generates an inactive CLV3 probe; biological activity is then reinstated upon light-triggered cleavage of the NVOC group. As before, the TAMRA fluorophore was linked to the AA2 position, giving fluorescently labeled, photoactivatable NVOC-CLV3-TAMRA (**8**) (Figure 6A).

**Figure 6:**
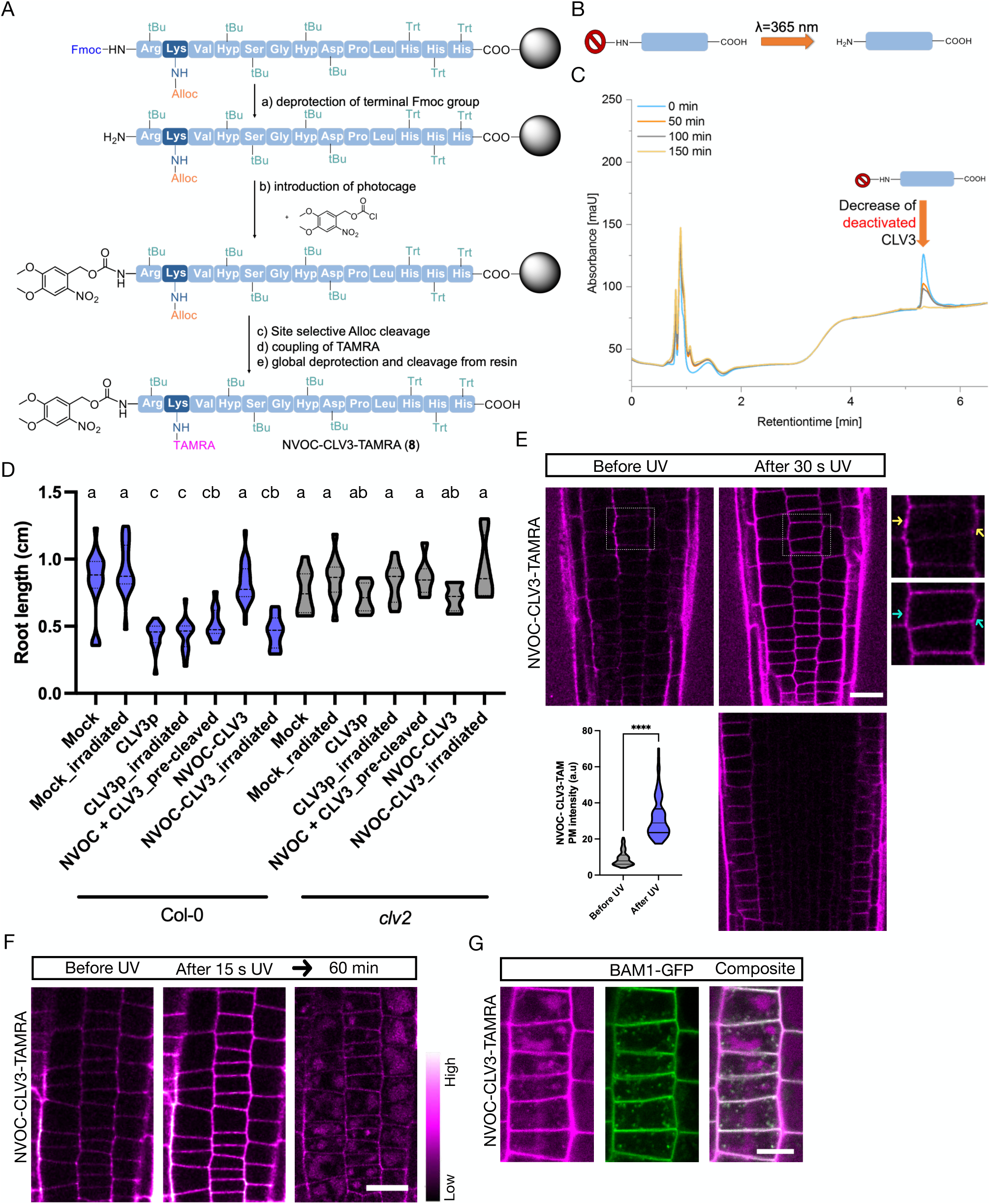
Synthesis and activation of photocaged peptide by UV radiation. A) Solid phase peptide synthesis of NVOC-CLV3-TAMRA using preloaded Fmoc-His(Trt)-Resin and Fmoc-protecting group (PG) standard protocol. Deprotection of N-Terminal Fmoc-PG (a) using 25 Vol% piperidin in DMF for 5-, 15- and 20min. Coupling photocage (b) using 5 eq. NVOC-Cl and 10 eq DIPEA in DMF/DCM 1:1 for 4h. Selective alloc deprotection (c) of lysin under reductive conditions with tetrakis(triphenylphosphine) palladium (0) catalyst and 10 eq. dimethylbarbituric acid in DCM two times for 45 min. Conjugation of TAMRA-N-hydroxysuccinimid (d) at free Lysine by adding 10 eq. DIPEA in DMF for 16h. Cleavage of the final structure (e) with 70 Vol% TFA and 30 Vol% TIPS. See Supplemental File 1 for further details on the synthesis and analytical data of photocaged peptides B) General scheme for peptide probe activation through 365 nm light-induced cleavage of N-Terminal NVOC photocage. C) Section of RP-HPLC chromatograms in which NVOC-CLV3 was irradiated with an UV medium-presure lamp including wavelength of 365 nm for a total duration of 150 minutes. After 150 minutes, a complete degradation of the NVOC-CLV3 signal was observed, indicating a complete cleavage of the NVOC-protecting group. See Supplemental File 1 for full Spectra. D) Violin plot representing root length analysis of 7-d old Col-0 and *clv2* seedlings 7-day treatment with mock, 200 nM CLV3p, 200 nM NVOC-CLV3 or pre-cleaved 200 nM NVOC-CLV3. The lines represent the median and the quartiles. n>9 roots; N=1. Two-way ANOVA with interaction, F test p<9.59e-12 ***. Statistical grouping was calculated by Tukey’s HSD test (**α**= 0.001). Groups sharing the same letter are not significantly different. E-G) Representative confocal images of root meristem epidermal cells after treatment with 1 µM NVOC-CLV3-TAMRA. Line used: *pBAM1::BAM1-GFP; bam1-3*. E) PM binding of NVOC-CLV3-TAMRA before and after photoactivation by UV exposure for 30s. Image of the epidermal layer (top). Image of the root centre with all layers visible (bottom). The inset shows the zoomed in cells with CLV3-TAMRA in the apoplast around the cells (yellow arrows) or at the PM (blue arrows). The violin plot represents the CLV3-TAMRA mean fluorescence intensity of the PM from 12-26 cells/root. n=4; N=6. Two-sided t test. For After > Before p<n0.0001. F) Vacuolar localization of photoactivated NVOC-CLV3-TAMRA 60 min after 15 s UV exposure. n=5; N=2. G) LE localization of photoactivated NVOC-CLV3-TAMRA 20 min after 15 s UV exposure. n=2; N=3. Scale bar: E, F – 20 µm; G – 10 µm.

Peptide synthesis was performed following previously discussed solid phase peptide protocols. Coupling of NVOC was performed on solid phase using NVOC chloride and coupling to the N-terminus of the terminal amino acid after Fmoc release. After installation of the photocage, the TAMRA fluorophore was coupled as previously described. After cleavage from the resin the crude peptide was purified by chromatography to give NVOC-CLV3-TAMRA with <90% relative purity (Data S1 and Data S3). First, release of the NVOC group upon light irradiation in solution was evaluated. Therefore, the NVOC-CLV3 peptide was exposed to UV light at 365nm and analyzed by RP-HPLC-MS-measurements at different time points. After 150 min, full deprotection was observed. Importantly, no decomposition or fragmentation of the deprotected peptide was found (Figure 6C, full spectra Data S4).

We first tested the efficiency of NVOC cleavage from the peptide and the bioactivity of the NVOC-cleaved CLV3. We performed root length assays on whole seedlings that were grown in medium containing NVOC-CLV3 or CLV3p or mock, that were either (*i*) irradiated with UV for 120min or (*ii*) not irradiated. UV irradiation of plates with NVOC-CLV3 resulted in shorter roots than controls, equivalent to CLV3p treated roots. This indicated that the NVOC group was cleaved off, thereby releasing CLV3 that inhibited meristem development. *clv2* mutants were resistant to NVOC-CLV3 with or without irradiation, indicating that the short roots were indeed due to activation of the photo-caged CLV3 peptide and signaling, and not a direct consequence of UV irradiation (Figure 6D). Roots protected from UV exposure developed similar to mock treated negative controls, showing that the NVOC group remained attached, which rendered the peptide inactive (Figure 6D). Additionally, the roots grown in medium with pre-cleaved peptide (NVOC+CLV3) also had shorter roots, proving that photoactivation of the photocaged peptide by UV is a reproducible technique and that NVOC in the medium does not interfere with the bioactivity of the peptide or plant development.

We then tested if the photocaged peptide can be rapidly photoactivated in the target tissue. To visualize peptide uncaging and perception, we photoactivated the NVOC-CLV3-TAMRA peptide (i) by UV irradiation at 365nm, which is the maximum absorption wavelength for NVOC cleavage(Schaper *et al*., 2009), or (ii) irradiation of targeted cells using the 405nm laser of a confocal microscope. 405nm is at the lower end of the absorption spectrum (Figure 6C), and might be less efficient than UV-light. NVOC-CLV3-Tamra was predominantly localized to the apoplastic space before photoactivation due to the blocked N-terminus. After 30s of UV irradiation using the UV lamp of the confocal microscope, we observed PM localization of the deprotected CLV3-TAMRA (Figure 6E - top). Notably, fluorescence was localized to the epidermal layer of the root (Figure 6E - bottom). 20 minutes after photoactivation, we observed CLV3-TAMRA in endosomal compartments, and subsequently also vacuolar localization (Figure 6F,G). Irradiation using the 405nm laser resulted in minor activation and significantly less PM localized fluorescent signal (Figure S4A).

We have successfully created photoactivable CLE peptide tools: NVOC-CLV3 and NVOC-CLV3-TAMRA, that can be activated simply by very short exposure to low energy UV radiation in target cells or tissue in vivo. The released bioactive peptide can be used to study rapid down-stream signaling responses or cell/tissue specific effects.

## 3. Discussion

Synthetic chemistry has aided in generating ligands to dissect ligand-receptor pairing, receptor activation and down-stream signaling. Such chemical tools have also been used to label phytohormones and peptides to probe their interaction with receptor kinases and subsequent signaling.(Sharma and Russinova, 2018) In this study we have synthesized a set of CLV3 probes with different modifications to study and decipher diverse aspects of CLE-RLK signaling.

Following the identification of CLE molecules, synthesis of the active CLE peptide enabled rapid elucidation of their effects and downstream signaling effectors through in vivo and in vitro bioassays.(Blümke *et al*., 2021; Breda *et al*., 2019; Breiden *et al*., 2021; Fiers *et al*., 2005; Ito *et al*., 2006; Qian *et al*., 2018; Stahl *et al*., 2009; Takahashi *et al*., 2018) Synthetic approaches have also provided a faster alternative to mutagenesis in creating peptide variants. Replacing individual or a combination of residues, and swapping of residues between peptides have facilitated breakthroughs in dissecting their affinities, specificities and recognition mechanism for individual RLKs.(Ito *et al*., 2006; Li *et al*., 2017; Roman *et al*., 2022; Song *et al*., 2012; Zhang *et al*., 2016a) However, most of those studies were performed in vitro. An alternative assay that allowed differentiation of affinities for RLKs was photoaffinity labeling (PAL), and binding of CLE9 and CLV3 peptides to several RLKs were resolved using this technique.(Ogawa *et al*., 2008; Shinohara and Matsubayashi, 2013, 2015) However, PAL is an expensive technique, requires elaborate protocols and does spatially resolve the ligand-receptor interactions within a tissue.(Sharma and Russinova, 2018) We emphasize here the advantages of fluorescently labeled peptides, which exhibit non-covalent reversible binding and native endocytic sub-cellular trafficking interactions. Attaching a fluorophore while maintaining biological activity is challenging, since CLE peptides are only 12/13 aa long, and modification of residues can change recognition and binding to cognate RLKs. Ogawa et al. had previously synthesized modified CLV3 peptides replacing L10 and T2, which still effectively bound the CLV1 ectodomain. Based on these in vitro experiments, we modified the second CLV3 residue to lysine (T2K) and added the fluorophore TAMRA.

Nevertheless, the second residue variation between CLE41 and −42 is enough to significantly alter their binding affinity to TDR.(Zhang *et al*., 2016a) Therefore, it is crucial to test both the bioactivity and specificity of the synthesized peptide tools. We observed that the peptide was indeed bioactive but showed reduced activity, which is likely caused by the attached TAMRA fluorophore. However, the peptide showed no significant decrease in affinity to CLV1.clade receptors, supporting the notion that CLV3-TAMRA peptide is functional. Furthermore, we utilized the probe to test for cross-clade interaction to the phylogenetically distant PEPR clade. We observed no binding affinity of CLV3-TAMRA to PEPR1. Importantly, pep1-TAMRA also showed no affinity to BAM1, a member of the CLV1-clade. This demonstrates that, in vivo, there is minimal cross-interaction of peptides from distinct families with different RLK clades. Gutiérrez-Alanís et al previously reported that CLE14 interacts with PEPR2. However, this interaction was only tested through a Bimolecular Fluorescence Complementation (BiFC) assay of CLE14-nYFP with PEPR2-cYFP, and it is likely that attaching a large fluorophore deactivated the CLE14-peptide.

Previous studies illustrated the importance of terminal residues for peptide function. Addition of an arginine group to the C-terminus significantly reduced CLE41 activity and its binding to TDR.(Ito *et al*., 2006; Zhang *et al*., 2016a) Similarly, unprocessed CLE19 proprotein with an extra C-terminal arginine exhibited diminished activity in vivo.(Tamaki *et al*., 2013) Alanine scanning experiments by Song et al. demonstrated that the N-terminal arginine residue of CLV3 is vital for its function, and addition of an extra tyrosine residue to the N-terminus of CLV3 or CLE45 rendered these peptides inactive.(Hazak *et al*., 2017; Song *et al*., 2012) Interestingly, we observed that a simple acetylation of the N-terminus was enough to deactivate CLV3. Based on this finding, we were able to manipulate the peptide N-terminus by attaching a photocleavable group that temporarily deactivates CLV3 and can be cleaved off efficiently with light at 365-380nm. This enables to track rapid or time-sensitive cell biological effects, and to elucidate tissue specific effects with spatial and temporal control. For example, rapid changes in sub-cellular localization of a protein of interest within seconds after CLE perception can be observed microscopically in vivo. Alternatively, the peptide can be activated in target cells and subsequent non-cell autonomous effects can be followed in adjacent cells or distant tissues. The rapid NVOC cleavage observed under the microscope, in comparison to in vitro experiments, can be attributed to several factors.

Firstly, differences in the light sources employed, such as lasers versus UV lamps, can lead to variations in energy output, potentially influencing the rate of cleavage. It is also crucial to note that the fluorescence observed under the microscope after 30 seconds does not necessarily indicate complete NVOC protecting group cleavage. In contrast, in vitro experiments involving spectral methods (e.g., absorbance and RP-HPLC spectra) offer a more quantitative assessment of cleavage completeness. The microscopic method confirms the occurrence of cleavage but does not provide the same level of quantitative information.

## Acknowledgement

We would like to thank Prof. Eugenia Russinova for sharing resources. We would like to acknowledge the Center for Advanced Imaging (CAi) at Heinrich-Heine-University Düsseldorf for providing access to the Zeiss LSM 880 Airyscan Fast funded by DFG-INST 208/746-1. LSM900 was funded through the ERC Synergy grant, Sympore. M.N and N.J were supported by Deutsche Forschungsgemeinschaft (DFG, German Research Foundation) through grants within the collaborative research center (CRC) 1208, B04 and A11 awarded to R.S and L.H, respectively.

## Author Contribution

Conceptualization: M.N, N.J, R.S, L.H; Methodology: M.N, N.J, E.B; Validation: M.N, N.J; Formal analysis: M.N, N.J; Investigation: M.N, N.J, E.B, F.K, F.B; Resources: R.S, L.H; Writing – Original draft: M.N, N.J; Writing – Review & Editing: M.N, N.J, R.S, L.H; Visualization: M.N, N.J; Supervision and Funding Acquisition: L.H, R.S. L.H. bears responsibility for synthetic chemistry and molecular characterization, R.S. for plant biology.

## Materials and methods

### Chemicals and Materials

Acetonitrile (≥99 %), dichlormethane (≥99 %) and acetic acid (≥99 %) were purchased from Merck. Acetic anhydride (99%) was purchased from VWR chemicals. Diethyl ether (stabilized with BHT, ≥99%) was purchased from Honeywell. 1,3-Dimethylbarbituric acid and Lithium chloride was purchased from Carl Roth. Piperidine (99%), tetrakis (triphenylphosphine) palladium (0), trifluoro acetic acid (99%) were purchased from Acros Organics. *N*,*N*-Dimethylformamide (DMF; for peptide synthesis) triisopropylsilane (TIPS, 99%), *N*,*N*-diisopropylethylamine (DIPEA, ≥99%), benzotriazol-1-yl-oxytripyrrolidinophosphonium hexafluorophosphate (PyBOP, 99%) was purchased from Fluorochem. 6-Carboxy-tetramethylrhodamine succinimidyl ester (TAMRA-N-hydroxysuccinimid) was purchased from Carbosynth. Dichlormethane (DCM, ≥99 %), 4,5-Dimethoxy-2-nitrobenzyl chloro formate (NVOC-Cl, 97%) were purchased from Sigma Aldrich. Fmoc-Hyp(tBu)-OH (99 %) was purchased from BLDpharm. Fmoc-Leu-OH (98 %) was purchased from Carbolution Chemicals. Boc-Arg(Pbf)-OH, Fmoc-L-Arg(Boc)2-OH, Fmoc-L-Arg(Pbf)-OH, Fmoc-L-Asp(OtBu)-OH, Fmoc-Gly-OH, Fmoc-L-Pro-OH, Fmoc-L-Ser(tBu)-OH, Fmoc-L-Thr(tBu)-OH und Fmoc-L-Val-OH were purchased from Iris Biotech with purities of 98,0%. Fmoc-His(Trt)-OH (98,0%), Fmoc-Lys(Boc)-OH (98,0%) und Fmoc-Pro-OH (99,0%) was purchased from Merck. H-L-His(Trt)-2CT resin (Loading 0,63 mmol/g) was purchased from Rapp Polymer. For TFA removal AG® 1-X8 resin from Bio-Rad Laboratories was used. Strata C18-E column (1g/ 6ml) was purchased from Phenomenex. Centrifugal Concentrator Vivaspin20 (1000 MWCO, 20ml) was purchased from Satorius.

### Plant materials

#### Arabidopsis thaliana ecotype

Col-0 was used as WT and generally as the background for the mutant and the transgenic lines, unless specified. *crn* and *clv2* mutant lines have been described as *crn-10* (CRISPR-Cas9-derived)(Nimchuk, 2017) and *clv2-gabi (GK-686A09)*(Pallakies and Simon, 2014), respectively. *crn-10* was verified using dCAPS strategy which involves amplifying the gene with the oligomers 5’-GTAGAAGCAGCAATGAAGCAAAGAAGAAGGTG-3’ and 5’-GTGTAGATGATGTTGAAGTT GTGGATAAGTG-3’ followed by HphI digestion. *bam1* single and *bam1;bam2* double mutants are CRISPR-Cas9-derived(Fan *et al*., 2021). The transgenic lines used in the study are: p35s::ARA7-GFP(Dettmer *et al*., 2006; Ueda *et al*., 2001), XVE>>AXL2(Adamowski *et al*., 2018), pBAM1::BAM1-GFP;*bam1-3*(Schlegel *et al*., 2021), *pUBQ10::VAMP711-YFP*(Geldner *et al*., 2009), pCLC2::CLC2-GFP (ecotype: Wassilewskija)(Konopka *et al*., 2008), pCLV1::CLV1-2xGFP; *clv1-11*(Nimchuk *et al*., 2011), RPS5::PEPR1-GFP; *pepr1; pepr2*(Ortiz-Morea *et al*., 2016).

### Macromolecule synthesis

#### General procedure for solid phase synthesis

All peptides were synthesized on solid phase using an automated synthesizer (Activotec P11). The standard Fmoc protocol was used. Therefore, Fmoc protected amino acids were used for the synthesis. H-L-His(Trt)-2CT resin (Loading 0,63 mmol/g) was used and the structures were synthesized by repetitive Fmoc cleavage and amide coupling. For Fmoc cleavage, the resin was treated three times with 25 Vol% piperidine in DMF for 10, 15 and 20 min. Afterwards the resin was washed 10 times with DMF. For coupling, the resin was treated with a solution of 5 eq. Fmoc amino acid, 5 eq. PyBOP and 10 eq. DIPEA in DMF for 1 h. Afterwards the resin was washed 10 times with DMF. After assembly of the full sequence the resin was washed 3 times with DCM. Thereafter, the modifications of the peptide described in detail below were performed. For final cleavage the resin was treated with 70 Vol% TFA, 30 Vol% TIPS for 1 h. The peptides were precipitated in diethyl ether, lyophilized and then TFA removal and further purification was perfomed.

#### Alloc deprotection and TAMRA labeling

After finishing the sequence the alloc deprotected of lysine on the resin was performed using 10 eq. 1,3 dimethylbarbituric acid and a spatula tip of tetrakis (triphenylphosphine) palladium (0) in DCM (1ml per 100 mg resin). First 1,3 dimethylbarbituric acid was dissolved DCM and flushed with argon for 2 min. The palladium catalyst was added and flushed with argon for 2 minutes. The solution was then incubated into the syringe for 45 min. Afterwards the resin was washed 10 times with DCM, 10 times with 0.2 M DIPEA in DMF and 10 times with DMF. The whole step was repeated once. After lysine was deprotected under reductive conditions, TAMRA conjugation took place. For this purpose, 1.3 eq. TAMRA-N-hydroxysuccinimid was dissolved in 4 ml of DMF and then 10 eq. of DIPEA was added. The solution was incubated in the syringe for 16 h. The resin was then washed with DMF 10 times and then alternately washed 3 times with DCM, 3 times with methanol until the solution becomes clear.

#### Acetylation

Acetic anhydride (1ml per 70 mg resin) was incubated to the syringe for 15 min 2 times. Afterwards the resin was washed 10 times with DMF and 10 times with DCM.

#### NVOC-Coupling

NVOC-Cl was dissolved in a 50:50 Vol% DCM/DMF mixture (1 ml per 100 mg resin). Then 10 eq. DIPEA was added and the mixture incubated in the solid phase syringe for 16 h. Afterwards the resin was washed 10 times with DMF and 10 times with DCM.

#### TFA removal

TFA removal was performed with a AG® 1-X8, quaternary ammonium, 100–200 mesh, acetate form resin. For 100 mg sample, 500 mg of the ion exchange resin was used. The resin was activated by washing 3 times with 10 ml of 1.6 M acetic acid solution, followed by 3 times with 10 ml of 0.16 M acetic acid solution. 100 mg sample were dissolved in 2 mL Milli-Q® and the solution was loaded to the resin into the syringe. The syringe was shaken for 1h. The supernatant was recovered, and the resin was washed 3 times with few Milli-Q®. The liquid phases were lyophilized to obtain the crude product as a white or pink solid (in case of TAMRA labelled peptides).

#### Vivaspin

Samples were separated from non-bound material using a Centrifugal Concentrator Vivaspin20 with 1000 MWCO. 5 times 10 ml with Milli-Q®, 5 times 10 ml 2 M LiCl solution, 5 times 10 ml Milli-Q® with 4400 rpm for 15 min each using an Hereaus Megafuge 8R.

#### Column purification

For NVOC-CLV3-TAMRA a further purification step was performed. A Strata C18-E column from Phenomenex (1g/ 6ml) was used to separate the samples from other incorrect sequences. First the column was conditioned with 6 ml acetonitrile and equilibrated with 6 ml Milli-Q®. 100 mg sample were dissolved in 1-2 ml of Milli-Q® and added to the column. Then the sample was washed with acetonitrile/Milli-Q® gradients from 5-30% each 3-6 ml on the column.

### Reversed Phase-High Pressure Liquid Chromatography (RP-HPLC) Electron Spray Ionization-Mass Spectrometry (ESI-MS)

RP-HPLC-MS spectra were performed on an Agilent Technologies 1260 Infinity Instrument in combination with a 6120-quadrupole mass spectrometer. The instrument has a wavelength detector (VWD1 A) that measures the absorbance at 214 nm. The mass spectrometer generates ions using the electrospray method, which is used at a Charge-to-mass ratio detected between 200 to 2000. The separation of the sample was performed on the MZ-AquaPerfect C18 column (3.0 x 50 mm, 3 μm) at 25 °C. As mobile Phase the following mixtures were used: A) H2O/acetonitrile (95:5 Vol.%) with 0.1 Vol.% Formic acid and B) H_2_O/acetonitrile (5:95 Vol.%) with 0.1 Vol.% formic acid. The flow rate was 0.4 ml/min. A linear gradient from 100%A/ 0%B to 0% A /100%B with a total measurement time of 17 min.

### Ultra-High Resolution - Mass Spectrometry (UHR-MS)

UHR-MS measurements were performed with a Bruker UHR-QTOF maXis 4G instrument with a direct inlet via syringe pump, an ESI source and a quadrupole followed by a Time of Flight (QTOF) mass analyzer

### MALDI-TOF-MS

MALDI-TOF-MS spectra were recorded on an UltrafleXtreme instrument from the company Bruker Daltonik. The concentration of the sample was 1 mg/ml. The ratio to the α-cyano-4-hydroxycinnamic acid (HCCA) matrix was 1:10.

### Lyophilization

The final oligomers were lyophilized with an Alpha 1-4 LD plus instrument from Martin Christ Freeze Dryers GmbH. The drying method was set to −40 °C and 0.1 mbar

### Seed sterilization and plant growth conditions

The seeds were sterilized using chloric gas (1 h in a desiccator after mixing 50 ml of 13% w/v sodium hypochlorite with 4 ml 37% HCl) and were sown on Murashige and Skoog (MS) medium containing ½ 0.22% w/v MS salts with B5 vitamins, 1% w/v sucrose, 0.05% w/v 2-(N-morpholino) ethanesulfonic acid (MES) and 12 g/l plant agar, adjusted to pH 5.7 with KOH. The seedlings for root experiments were grown in phytocabinets (poly klima; model: M4Z-TDL+rF) for 4 or 5 days vertically with continuous light at 21°C. Later, they were transferred to soil and grown in phytochambers for 5 to 6 weeks under 16 h light/8 h dark condition to experiment on the shoot apical meristem. The light spectrum for plant growth condition spans the wavelengths from UV to far red with very minimal irradiance of 1-5 mW/m^2^/nm for wavelength <400 nm, and 11-170 mW/m^2^/nm for photosynthetically active radiation (PAR) of 400-700 nm range.

### Treatment and imaging conditions

#### Peptide treatment

The pep1 (synthesized by Davids Biotechnologie), pep1-TAMRA peptide (obtained personally from Prof. Russinova), mock controls and peptides except CLV3p (synthesized by Davids Biotechnologie) were dissolved in water to make stocks and subsequently added to the MS medium to reach the indicated working concentration. CLV3p was dissolved in peptide buffer of pH 6 (mixture of 87.7 ml of 0.2 M potassium phosphate, mono-potassium salt and 12.3 ml of 0.2 M potassium phosphate, di-potassium salt in 100 ml buffer).

#### Chemical treatment

4-d-old seedlings of XVE>>AXL2 line were transferred to MS agar plates containing 10 µM estradiol (10mM stock dissolved in DMSO) for 24-hour AXL2 induction. For uninduced control condition, seedlings were transferred to MS agar plates containing the solvent DMSO. Subsequent mock or peptide treatments in AXL2 induced seedlings were also made in MS agar plates containing either 10 µM estradiol or DMSO.

BFA treatments on seedlings were done in GM liquid medium containing 50 µM BFA together with 1 µM peptide or the controls for 50 min.

#### Sample handling and imaging

In vivo live fluorescence microscopy was performed at the CLSM systems Zeiss LSM 880 and Zeiss LSM 900 employing C-Apochromat 40x/1.20 water objectives. For root length assays, the MS agar plates were scanned using CanoScan 9000F with 600 dpi resolution.

For root meristem imaging, 4- or 5-day-old seedlings were incubated for the indicated time in either MS medium (with or without agar) containing mock or peptide. For photoactivation of the NVOC-CLV3-TAMRA peptide, the seedlings were mounted with MS medium containing the peptide and directly exposed to UV light (UV lamp – X-Cite XYLIS/ model XT720L) with DAPI filter.

For shoot apical meristem imaging, the entire inflorescence of a 5 or 6-week-old plants was submerged in water containing mock or peptide, 0.01% Tween 20 and 0.1% dimethyl sulphoxide (DMSO). The inflorescence was then cut off then mounted onto the slide over a double-sided adhesive tape. It was then dissected to expose the apical meristem. For CLV1 localization assay, the inflorescences were treated for 30 s and imaged after 30 min. For CLV3-TAMRA binding assay, the inflorescences were treated for 1 min, washed thrice and imaged immediately.

For root length assays, the seeds were directly sown on MS agar plates containing mock or peptide. For the root length assays with NVOC-CLV3, 3 days after germination, one set of plates with peptides or mock were exposed to UV black light (OUSIDE) for 3 h at a distance of 25 cm while the control set were kept unexposed. The plates were covered in yellow foil during growth to filter off light under 500 nm in order to avoid NVOC cleavage before and after UV radiation. After the indicated number of days, the plates were scanned and the length of the roots were analyzed. For pre-cleaving the NVOC-peptide, a solution of 0.5mg/ml was prepared in Milli-Q®. It was filled in a cuvette and radiated with UV black light for 3 h. The pre-radiated NVOC peptide were examined with UV-Spectra, RP-HPLC-MS and furthermore used for the root length assays.

### Testing the degradation of NVOC group

The samples were irradiated from a distance of 25 cm under UV medium pressure lamp (Heraeus Noblelight) fitted with an Hg lamp that emit light in the 250-600 nm range, or 50 W UV black light (OUSIDE) fitted with COB LED chip that emits light in the 395-400 nm range.

### Image analysis

Root length analysis and the PM and cytosolic intensity analysis were made using Fiji ImageJ software tools. Number of SAMs with vacuolar CLV1 localization and the number of root meristem cells with PM PEPR1 were recorded by visual inspection. The plots were made using GraphPad Prism 9 and Origins.

### Statistical analysis

All the statistical tests on intensity measurements were made using GraphPad Prism 9. Statistical tests for root length analysis were made using R version 4.3.1. ‘n’ indicates biological replicates and ‘N’ indicates technical replicates.

## Supplementary file 1

