## Supplemental figure 1 for "Macromolecular toolbox to elucidate CLE-RLK binding, signaling and downstream effects"

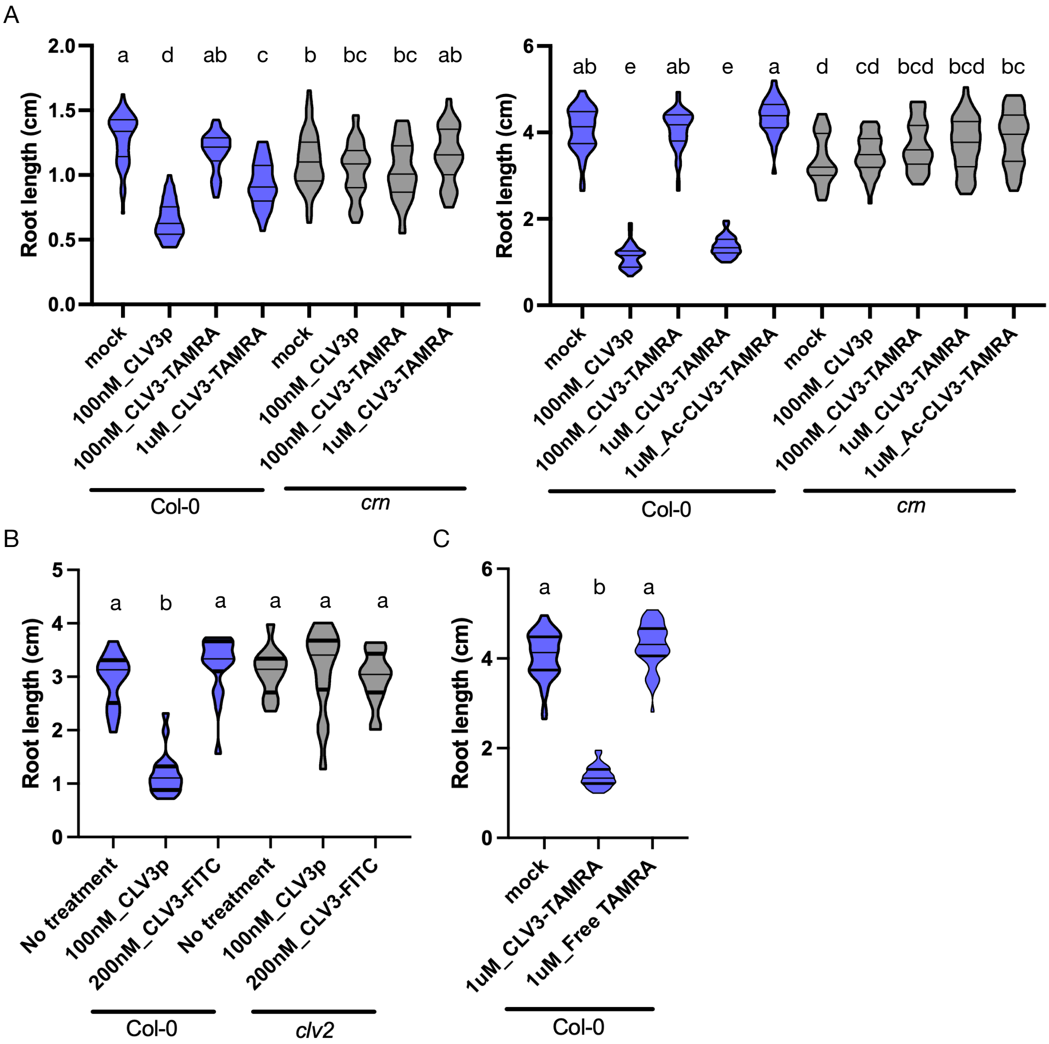


**Figure S1: Bioactivity of CLV3-TAMRA (Related to Figure 3)**

A – C) Violin plots representing root length analyses of Col-0 or *crn/clv2* mutant after respective treatments. The lines represent the median and the quartiles. Statistical grouping was calculated by Tukey's HSD test (**α**= 0.001). Groups sharing the same letter are not significantly different. A) After 5-day (left) and 10-d (right) treatment with mock, CLV3p, CLV3-TAMRA and Ac-CLV3-TAMRA. n≥ 33 roots; N=3. Two-way ANOVA with interaction, F test p<2e-16 ***. B) After 8-day treatment with mock, CLV3p and CLV3-FITC. For Col-0/mock > 200nM_CLV3-FITC p= n.s. n≥ 12 roots; N=1. Two-way ANOVA with interaction, F test, p<6.94e-13***. C) After 10-day treatment with mock, CLV3p and free TAMRA fluorophore. For Col-0/mock > Free TAMRA p= n.s. n≥ 46 roots; N=3. One-way ANOVA, F test, p<2e-16 ***.
