## Supplemental figure 2 for "Macromolecular toolbox to elucidate CLE-RLK binding, signaling and downstream effects"

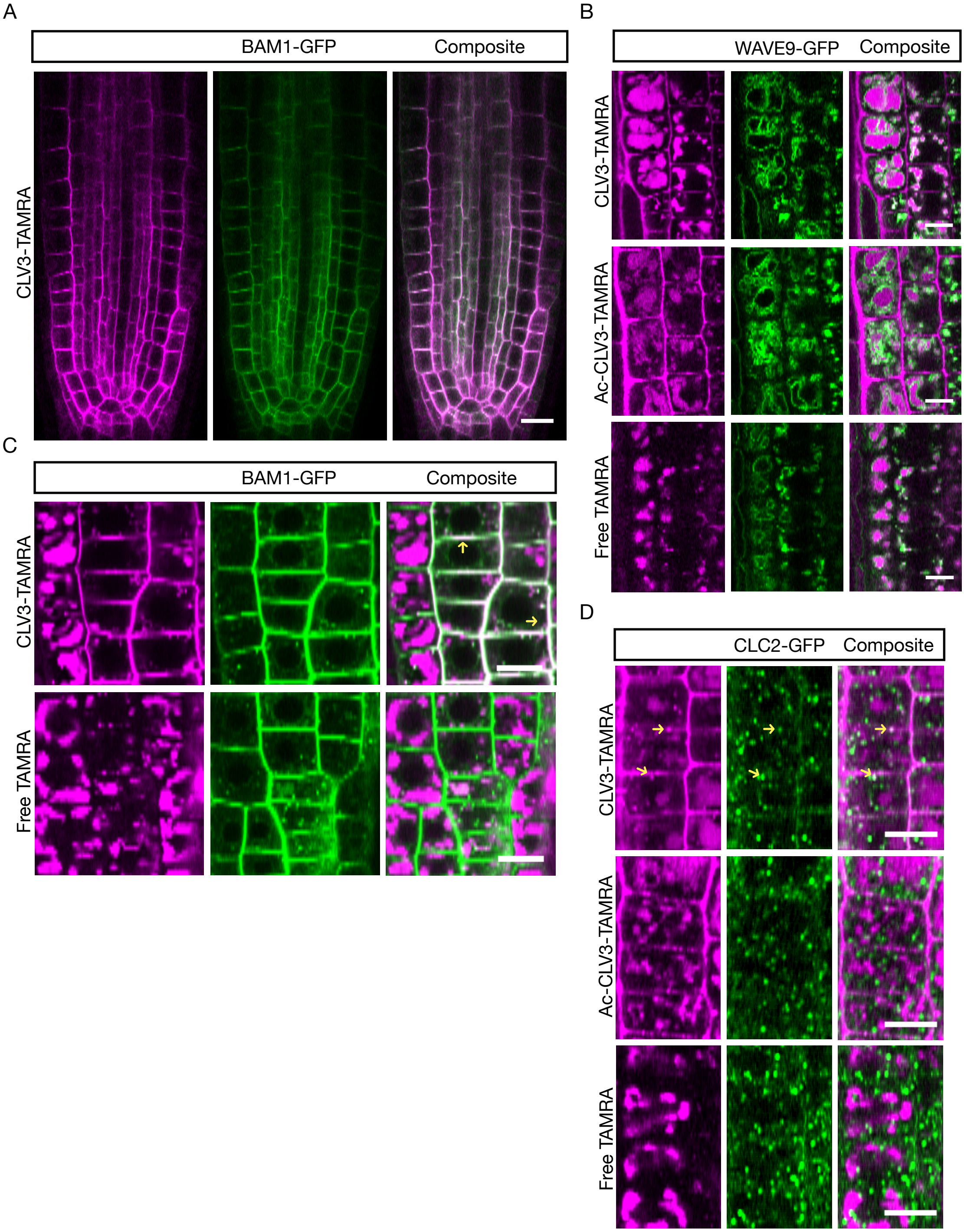


**Figure S2: Sub-cellular localization and trafficking of CLV3-TAMRA (Related to figure 4)**

A-D) Representative confocal images of root meristem epidermal cells after 1 $\mu$M treatment of the peptide CLV3-TAMRA, or the controls Ac-CLV3-TAMRA or free TAMRA fluorophore. Lines used: A, C – *pBAM1::BAM1-GFP; bam1-3*; B – WAVE9Y (*pUBQ10::VAMP711-YFP*); D – *pCLC2::CLC2-GFP*.
A) PM localization after 3 min treatment. n≥5; N=2. PM marked by BAM1-GFP. B) Vacuolar localization after 10 to 15 min post 15 min treatment. n≥5; N=2. Vacuoles marked by tonoplast localised protein VAMP711-GFP. C) PM localization after 60 min treatment. n≥5; N=2. BAM1-GFP marks the PM and the endosomes. The arrows indicate an example of PM and endosomal co-localization. D) EE localization 10 to 15 min post 5 min treatment. n≥5; N=2. The arrow indicates CLV3-TAMRA co-localized with the EE marker, CLC2-GFP. Note: In B-D, Ac-CLV3-TAMRA and Free TAMRA were imaged at higher laser intensity and gain than CLV3-TAMRA to observe their sub-cellular localization. Scale bar: A – 15 $\mu$m; B-D – 10 $\mu$m.
