## Supplemental figure 3 for "Macromolecular toolbox to elucidate CLE-RLK binding, signaling and downstream effects"

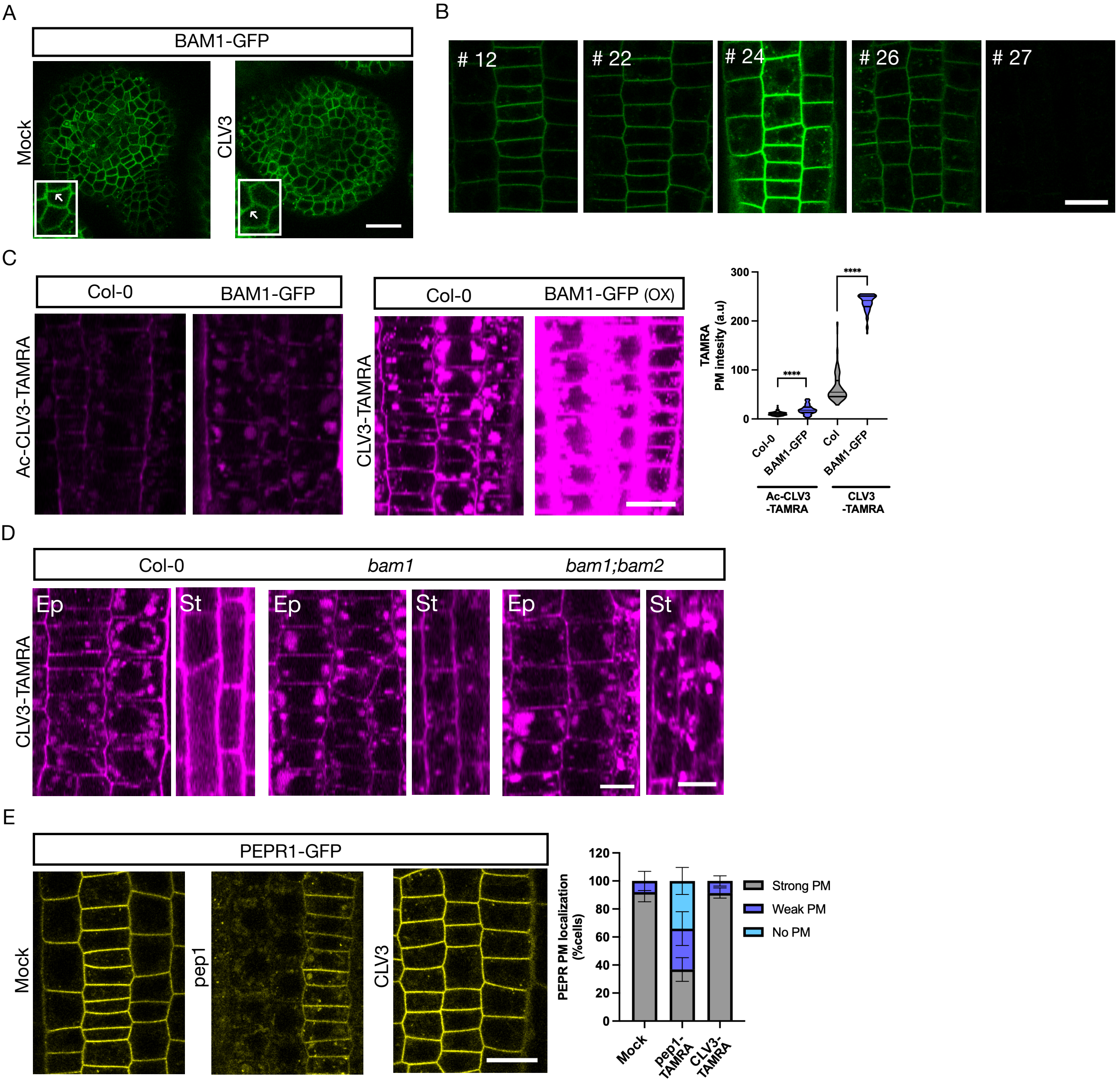


**Figure S3: Binding specificity of CLV3-TAMRA to RLKs (related to figure 5)**

A-E) Representative confocal images of SAM (A) or root meristem epidermal cells (B-E). Lines used: A, B – *pBAM1::BAM1-GFP; bam1-3*; C – Col-0 and *pBAM1::BAM1-GFP*; *bam1-3;* D – Col-0, *bam1 and bam1;bam2;* E – *pPEPR1::PEPR1-GFP*; *pepr1 pepr2*. A) BAM1 localization after 30 min to 2h of treatment with either mock or 100 nM CLV3.. N=6. The arrows indicate PM localised BAM1. B) BAM1-GFP PM level in independent lines. n≥5; N=1. C) CLV3-TAMRA PM signal after 1 $\mu$M, 15 min treatment in Col-0 or BAM1-GFP over-expression line. n=6; N=1. The violon plot represents Ac-CLV3-TAMRA or CLV3-TAMRA mean fluorescence intensity of the PM from 8-27 cells/root. The lines indicate the median and the quartiles. Two-sided t test. For Ac-CLV3-TAMRA/BAM1-GFP > Col-0 p<0.0001; For CLV3-TAMRA/BAM1-GFP > Col-0 p<0.0001. D) CLV3-TAMRA PM signal in epidermis (Ep) and stele (St) cell layers after 5-15 min post 5 min 1$\mu$M treatment. n=7; N=2. E) PEPR1 PM localization after 30 s pulse with mock, 100 nM pep1 or CLV3. The stacked bar graph represents relative number of cells with strong, weak or no PEPR at the PM. n=6; N=1. At least 58 cells were analyzed/root. Error bars indicate mean ± SD. Scale bar: A-C, E – 15$\mu$m; D – 10 $\mu$m (Ep), 7 $\mu$m (St).
