## Supplemental figure 4 for "Macromolecular toolbox to elucidate CLE-RLK binding, signaling and downstream effects"

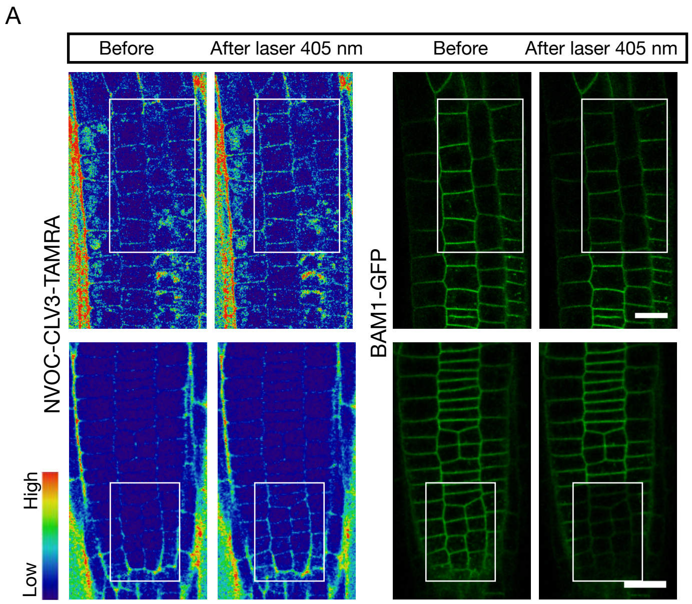


**Figure S4: Activation of photocaged peptide by 405nm laser irradiation (Related to figure 6)**

A) Representative confocal images of NVOC-CLV3-TAMRA (1$\mu$M) at the PM before and after photoactivation in target cells by 405nm laser (left). Corresponding BAM1-GFP PM intensity after laser activation in target cells (right). Line used: *pBAM1::BAM1-GFP; bam1-3*. n=8; N=3. Scale bar: 20$\mu$m.
